# Interindividual differences in predicting words versus sentence meaning: Explaining N400 amplitudes using large-scale neural network models

**DOI:** 10.1101/2025.06.03.657727

**Authors:** Milena Rabovsky, Alessandro Lopopolo, Daniel J. Schad

## Abstract

Prediction error, both at the level of sentence meaning and at the level of the next presented word, has been shown to successfully account for N400 amplitudes. Here we address the question of whether people differ in the representational level at which they implicitly predict upcoming language. To this end, we compute a measure of prediction error at the level of sentence meaning (magnitude of change in hidden layer activation, termed semantic update, in a neural network model of sentence comprehension, the Sentence Gestalt model) and a measure of prediction error at the level of the next presented word (surprisal from a next word prediction language model). When using both measures to predict N400 amplitudes during the reading of naturalistic texts, results showed that both measures significantly accounted for N400 amplitudes even when the other measure was controlled for. Most important for current purposes, both effects were significantly negatively correlated such that people with a reversed or weak surprisal effect showed the strongest influence of semantic update on N400 amplitudes, and random-effects model comparison showed that individuals differ in whether their N400 amplitudes are driven by semantic update only, by surprisal only, or by both, and that the most common model in the population was either semantic update or the combined model but clearly not the pure surprisal model. The current approach of combining large-scale models implementing different theoretical accounts with advanced model comparison techniques enables fine-grained investigations into the computational processes underlying N400 amplitudes, including interindividual differences.

## Introduction

Language processing is often assumed to be predictive, in the sense that readers and listeners routinely and probabilistically anticipate upcoming language input (e.g., Kuperberg & Jäger, 2016). One crucial piece of evidence supporting this assumption is the N400 component of the event-related brain potential (ERP), a negative component measured at centro-parietal electrode positions, whose amplitude gradually increases with decreasing fit or predictability of a given word in a given context (Kutas & Hillyard, 1980; see Kutas & Federmeier, 2011, for review). Even though the component’s relation to prediction and prediction error is now widely acknowledged, there is still an active debate about the specific cognitive mechanisms underlying this brain signal.

Recent years have seen a surge of computationally explicit theories on the functional basis of the N400. While many of them link the N400 to prediction error, they vary in terms of its assumed representational level (e.g., Rabovsky & McRae, 2014; Frank et al., 2015; Rabovsky et al., 2018; Fitz & Chang, 2019). Specifically, one important dimension along which these models differ concerns whether they link N400 amplitudes to prediction error at the word or sentence level.

Word-level prediction error has been often modeled by surprisal, that is the inverse probability of a given word in a given context. Surprisal from computational language models has been shown to predict N400 amplitudes during the reading of naturalistic texts (e.g., Frank et al., 2015; Heilbron et al., 2022; Michaelov et al., 2023). Prediction error at the sentence level has instead been modeled as the activation change in a hidden layer representation implicitly representing predicted sentence meaning (a measure called semantic update) within the Sentence Gestalt (SG) model, a neural network model of sentence comprehension (McClelland et al., 1989). Semantic update has been found to reproduce a broad range of empirically observed N400 effects, establishing a connection between this component and prediction error at the level of sentence (or more generally: message) meaning (Rabovsky et al., 2018).

Thus, prediction error both at the level of sentence meaning and at the level of the next presented word has been shown to successfully account for N400 amplitudes. What does this mean? Because these two forms of prediction error are positively correlated, it seems possible that only one of the measures constitutes the “real” cognitive correlate of the N400, and that the other measure explains variance in N400 amplitudes via its correlation with this “real” cognitive correlate. Alternatively, N400 amplitudes might reflect prediction error both at the word and at the sentence level. Here, we specifically investigate the possibility that people might differ in their level of implicit prediction, with some people more focused on predicting sentence meaning and others more focused on predicting the next upcoming word.

Until very recently, addressing such issues would have been difficult. This is because, apart from surprisal, which can be derived from computational language models including from deep learning0020models, other computationally explicit models of the N400, especially the cognitively motivated neural network models, were small-scale models trained on toy corpora and thus could only be related to the N400 in a qualitative way. To overcome these limitations and to be able to relate the model’s N400 correlate directly and quantitatively to EEG data collected from human participants presented with the same stimuli, recently a SG model has been trained on a large-scale corpus. Specifically, a SG model and a next word prediction language model that were as similar as possible in terms of architecture and number of parameters have been trained on the same large-scale language corpus (Sayeed et al., 2018) in order to enable a fair quantitative comparison between both measures of interest and thus between both theoretical accounts of the N400. Then semantic update (SU) from the SG model and surprisal from the language model were used to predict N400 amplitudes obtained during the reading of naturalistic text, that is text that does not contain any specific manipulations but rather just natural variations of expectancy as encountered in everyday life (using an existing EEG dataset that has previously been used to demonstrate the influence of surprisal on the N400; Frank et al., 2015). Results show that both semantic update and surprisal account for N400 amplitudes even when the other measure is controlled for (Lopopolo & Rabovsky, 2024).

Here, in order to investigate whether people differ in whether they implicitly focus on predicting word or sentence meaning, we use linear mixed-effects models to specifically investigate random effects correlations between the effects of semantic update and surprisal. Moreover, a novel and exciting approach to model comparison of neuro-cognitive models is to allow the true computational model and its parameters to differ between individuals, effectively treating the model as a random effect in a hierarchical variational Bayesian approach (Piray et al., 2018). Here, we use this random-effects Bayesian model comparison approach to study whether the true computational model generating N400 amplitudes differs between individuals.

To foreshadow our results, we find a negative random effects correlation between semantic update effects and surprisal effects, suggesting that for those people whose N400 amplitudes were most strongly predicted by semantic update, there was a weaker (and partly reversed) influence of surprisal. Moreover, random-effects model comparinformation based on PropBank information based on PropBank information based on PropBank ison shows that for a similar percentage of people N400 amplitudes are elicited by (i) only semantic update versus (ii) semantic update and surprisal combined, whereas for only few people only surprisal elicits N400 amplitudes. Thus, people may differ in their implicit prediction strategies with some people focusing more on expected sentence meaning and others focusing on both levels of words and sentence meaning at the same time, whereas only few people seem to focus purely on predicting the next word.

## Methods

### Model architecture

Model architectures are depicted in Figure 1. The **Sentence Gestalt (SG) model** maps sentences to representations of the described events (McClelland et al., 1989; see ‘Model tasks’ for details). It comprises two main components: an update network and a query network. The update network sequentially processes each incoming word to update the activation of the SG layer. This layer represents the meaning of the sentence after the presentation of each word, functioning as a combination of its previous activation and the activation induced by the new incoming word. The update network consists of an input layer, generating a vectorial representation w_t_ for each input word i_t_ in the incoming sentence. It also includes a recurrent Long Short-Term Memory (LSTM) layer that generates a SG representation sg_t_ as a function of w_t_ and the previous gestalt sg_t-1_. It maintains a hidden state that evolves as each new word is processed, enabling the model to retain information from earlier parts of the sequence. The query network extracts information about the event described by the sentence from the activation of the SG layer. The sentence comprehension mechanism is implemented in the update network, while the query network is primarily used for training. It is composed of a hidden state h_t_, generated by combining the SG vector sg_t_ and probe vectors p_i_. The output o_i_ of the query network is then generated from the hidden state h_t_.

**Figure 1.**
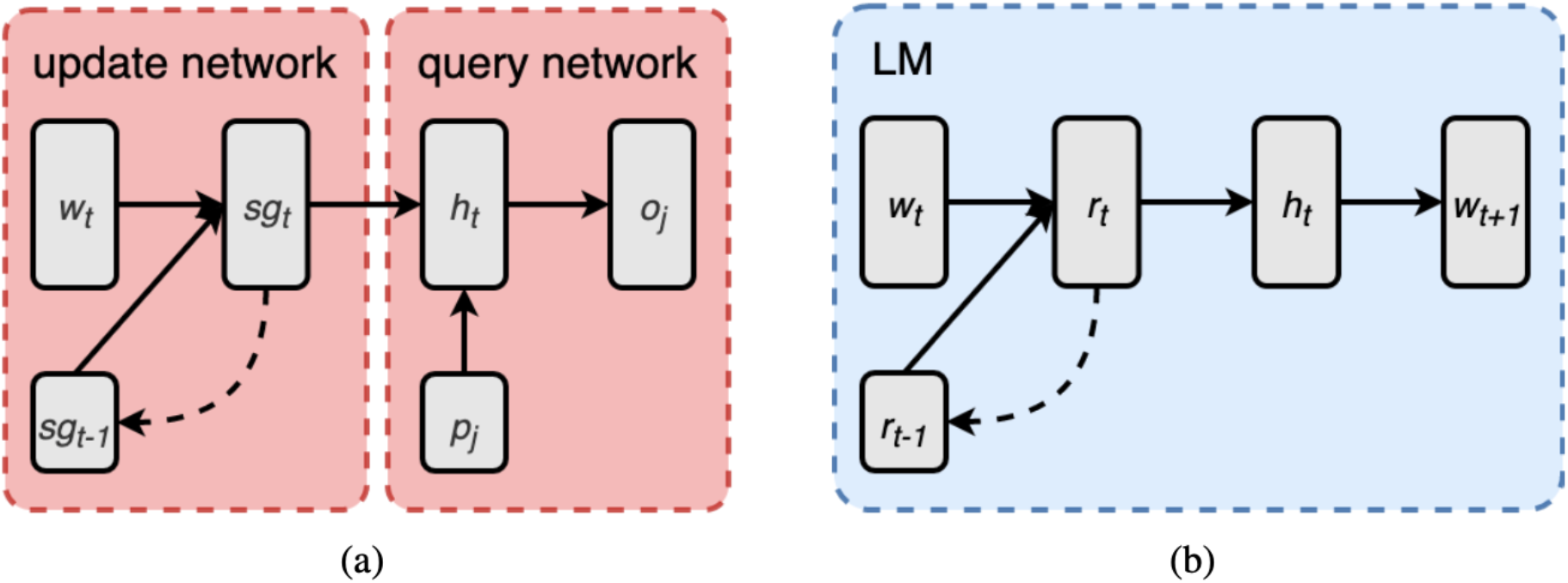
The architectures of (a) the Sentence Gestalt (SG) model and (b) the Language Model (LM).

The **next-word prediction language model (LM)** is designed to predict the probability distribution of the next word in a sequence given the context of preceding words. It also employs a recurrent neural network based on a LSTM architecture. The LM consists of an input layer, a recurrent layer, a feed-forward hidden layer and an output layer. The input layer encodes the word representations. The recurrent layer processes the sequential input and captures contextual information. The output layer produces a probability distribution over the vocabulary, indicating the likelihood of each word being the next in the sequence.

### Model tasks

The **task of the SG model** is to map a sentence to its corresponding event, defined as a list of role-filler pairs representing an action or state, its participants (e.g., agent, patient, recipient), and eventual modifiers. The event is implemented as a set of role-filler pairs represented as vectors o_i_, each formed by concatenating the feature representation of a word and a one-hot vector indicating the role of that word in the context of the sentence. For instance, the sentence ``the boy ate soup at lunch’’ consists of a sequence of six one-hot word representation vectors. Its event structure, however, contains four role-filler vectors representing each role of its event (agent, action, patient, time) with its corresponding concept filler (boy, eat, soup, lunch); see Lopopolo & Rabovsky (2024) for details.

During training, the model is presented with sentences fed word by word to the input layer. Every time a word is presented, the model is probed regarding the event described by the sentence. The model is probed for the complete event, even if the relevant information has not yet been presented at the input layer. Thus, during training the model’s connection weights are optimized for predicting the meaning of the complete sentence based on the information provided so far at the input layer and its experience of the statistical regularities in the training corpus. A probe consists of a vector p_i_ of the same size as a corresponding role-filler vector p, but with either the thematic role identifier zeroed (if probing for roles) or filler features zeroed (if probing for fillers). Responding to a probe involves completing the role-filler vector. When probed with either a thematic role or a filler, the model is expected to output the complete role-filler vector.

The **task of a next-word prediction LM** is to predict the most likely word to follow a given context. The training involves presenting the model with sentences one word at a time and adjusting its connection weights to optimize the accuracy of predicting the next word.

### Model-based variables

We compute two measures from our models: Semantic Update (SU) and surprisal. **Semantic Update (SU)** refers to the update of the SG model’s recurrent layer’s representations after the presentation of each word in a sentence. It is computed as the mean absolute difference between the activation of the SG layer before and after the presentation of a word. SU quantifies the amount of change induced by the new incoming word in the implicit predictive representation of sentence meaning. **Surprisal** instead is the negative log-probability of the new incoming word given the previous words: surp(w_t_)=-logP(w_t_|w_1:t-1_). It is computed from the probability distribution generated as output of the LM. Surprisal operates on lexical items, not on semantic representations.

### Training corpus

Both models were trained on the British National Corpus section of the Rollenwechsel-English (RW-eng) corpus (Sayeed et al., 2018). The RW-eng corpus is annotated with semantic role information based on PropBank roles (Palmer et al., 2015). The SG model is trained on mapping each RW-eng sentence to its PropBank-style event structure, while the LM is trained on predicting the next word in the sequence of each RW-eng sentence.

### EEG data

The electrophysiological recordings of the N400 were obtained from an EEG dataset provided by Frank et al. (2015), which consists of data collected from twenty-four participants (10 female, mean age 28.0 years, all right handed and native English speakers) while they were reading English sentences. The stimuli consist of 205 sentences from the UCL corpus of reading times (Frank et al., 2013), and originally from three little known novels. The sentences were presented in random order, word by word. The N400 amplitude for each participant and word token was defined as the average potential over a 300-500 ms time window after word onset at electrode sites in a centro-parietal region of interest. For further details regarding the stimuli see Frank et al. (2013), and concerning the EEG dataset, its stimulation paradigm and preprocessing see Frank et al. (2015).

## Results

Data and code for all analyses will be made available on OSF upon publication. We ran a linear mixed-effects model (LMM) to explain N400 amplitudes based on the fixed effects baseline EEG response, the semantic update (SU) of the SG model, and the surprisal calculated from a LM. As random effects, we incorporated correlated random intercepts and random slopes for the semantic update and the surprisal effects across subjects, as well as random intercepts across words. The lmer package (Bates et al., 2015) together with the lmerTest package (Kuznetsova, Brockhoff, & Christensen, 2017) were used to obtain p-values for the fixed effects.

As reported before (Lopopolo & Rabovsky, 2024), the results showed a strong effect of semantic update on N400 amplitudes (b = 0.213, SE = 0.035, df = 40.2, t = 6.012, p = 4e-7, standardized β = 0.055, 95% CI [0.036 0.074]. Moreover, there was a clear effect of surprisal (b = 0.163, SE = 0.056, df = 44.9, t = 2.902, p = .006, standardized β = 0.042, 95% CI [0.013 0.072]. These results show that both theoretical quantities – surprisal from a LM and semantic update from the SG model overall influence the size of the N400 even when the other quantity is taken into account. Here, we specifically looked at interindividual differences that were estimated by the linear mixed-effects model. First, the results showed interindividual variability for both effects: the semantic update effect on the N400 showed a standard deviation of 0.118 across participants, which with a mean of b = 0.213 suggests that the variation in the effect size was not too large. Indeed, Figure 2a (left panel) shows that the predicted effect size (i.e., individual regression coefficient predicted from the LMM) was positive in all 24 participants, suggesting that variation exists in how strongly semantic update influences the N400, but that the effect goes in the same direction for all participants. Concerning individual variability of the surprisal effect across participants, we found a by-participant standard deviation of 0.216, which is already larger, given the mean estimate of b = 0.163. Indeed, Figure 2a (right panel) shows that while in the majority of participants, high surprisal leads to a larger N400, there is a subset of 5 participants (shown in red) for which the effect flips sign, and becomes (at least qualitatively) negative, such that high surprisal is associated with a smaller N400. This interaction between surprisal and participant thus qualifies the interpretation of the fixed effect of surprisal, since surprisal effects cannot be concluded to be positive in all participants.

**Figure 2.**
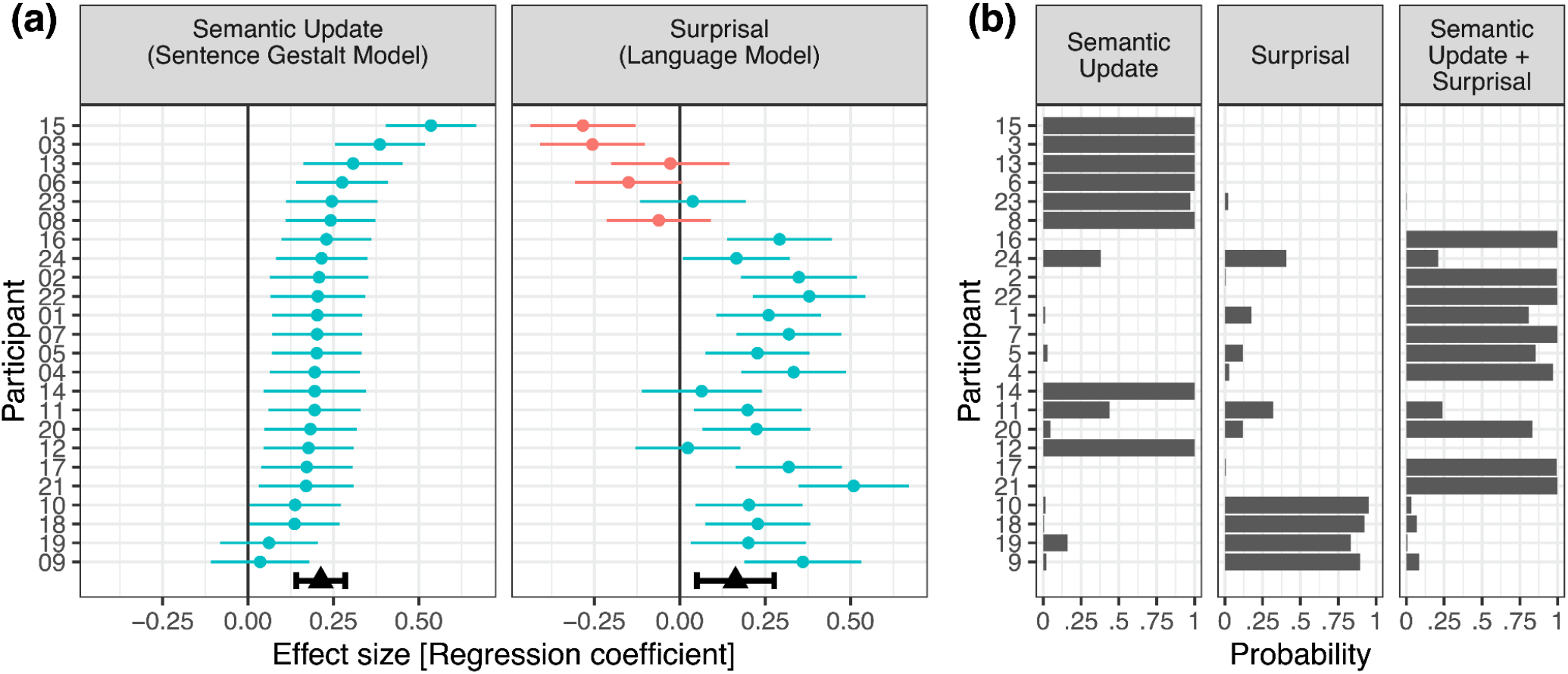
Interindividual differences in computations underlying N400 amplitudes. (a) Random slopes for semantic update and surprisal effects on N400 amplitudes. The predicted regression coefficient from a linear mixed-effects model is shown for each participant, error bars indicate 95% prediction intervals. Green color indicates participants with a positive regression coefficient, red color indicates participants with a negative regression coefficient. Black triangles and error bars at the bottom of the panels show fixed effects estimates with 95% CI. (b) Posterior model probabilities (i.e., responsibilities, that is the probabilities by which each participants’ data was generated by a given model) for each participant from a hierarchical Bayesian analysis, treating models and their parameters as random effects (Piray et al., 2018). Each of the three models is a linear model with the computational predictors semantic update / surprisal / both. (a+b) Participants are sorted by the size of their semantic update effect in the linear mixed-effects model.

Figure 2 orders the subjects by the size of their predicted semantic update effect on the N400, with large semantic update effects being shown at the top. Interestingly, it is visible that the subjects with a negative surprisal effect (shown in red) are all clustered at the top of Figure 2a (right panel), suggestive of a random effects correlation between semantic update effects and surprisal effects. Indeed, the random effects correlation as estimated by the LMM shows a negative correlation of r = -0.65 which is significantly smaller than zero, 95% CI [-1.00, -0.15], based on profiling. The resulting random effects correlation (as estimated from the LMM) is visualized in Figure 3. It confirms that individuals with a large surprisal effect show small to moderate semantic update effects, whereas in individuals with a negative surprisal effect, the semantic update effect is enhanced. We further followed up on this effect, and tested it using model comparison, by constraining all random effects correlations to zero, and then including the critical random effects correlation back into the model. The resulting model comparison again showed a significant effect, **Χ**^2^(1) = 5.98, p = .014. Also, AIC supported the more complex model (M0: AIC = 202855, M1: AIC = 202851, delta AIC = 4). To further statistically confirm this random effects correlation, we computed the semantic update and the surprisal effect for each participant separately, in a separate linear model of the N400 signal with predictors EEG baseline, semantic update, and surprisal. The resulting individual regression coefficients were significantly negatively correlated across subjects, with r = -0.47, t(22) = -2.485, p = 0.021, 95% CI [-0.73, -0.08].

**Figure 3.**
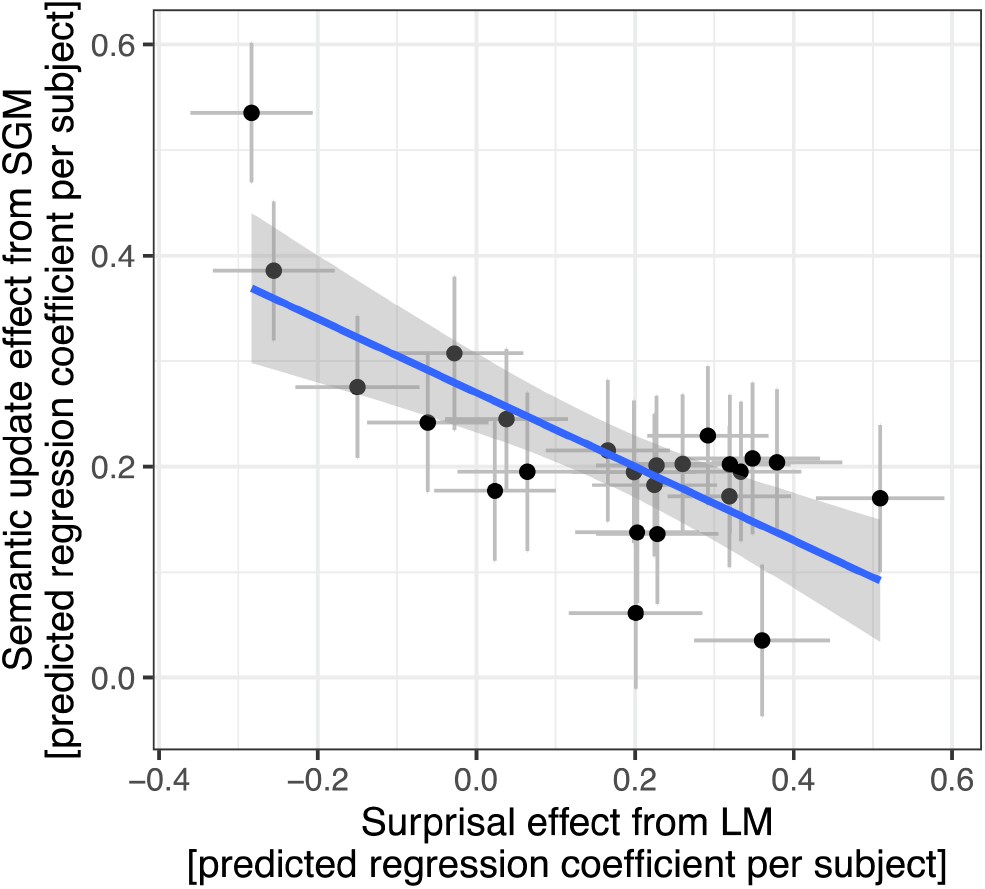
Random effects correlation between surprisal effects and semantic update effects. The semantic update and surprisal effects are predicted regression coefficients per participant from a linear mixed-effects model, with 95% prediction intervals. The line shows predictions from a linear (unweighted) regression model with 95% confidence bands.

The results from the linear mixed-effects model suggest that participants may differ in the computational processes that underlie N400 amplitudes, i.e., for some participants, N400 amplitudes may be elicited by both computations (semantic update and surprisal), whereas for other groups of participants, N400 amplitudes may be generated only by surprisal, or only by semantic update. Taking this result seriously, this amounts to assuming a random effect for models, i.e., that for different participants, different (computational) models may underlie N400 amplitudes. We next perform hierarchical Bayesian model selection and parameter estimation using a method that was recently developed to address such situations with random effects for models and their parameters (Piray et al., 2018). We allow for the possibility that for different participants, one of three different models underlies N400 amplitudes: (i) only semantic update, (ii) only surprisal, or (iii) a combination of semantic update and surprisal. All models contain baseline EEG amplitudes as control covariate. Regression coefficients for surprisal and semantic update were constrained to be positive using an exponential transform, i.e. negative regression coefficients were not taken as positive evidence for the respective model. Priors for model parameters were uncorrelated normal distributions with means of zero and standard deviations of one.

Results from this analysis show that subgroups of participants exist for which only semantic update, only surprisal, or a combination of both measures underlies N400 amplitudes. Figure 4 (left panel) shows that for the largest percentages of participants, either the combination of both models (42%) or the pure semantic update model (38%) generates N400 amplitudes, whereas for fewer participants the pure surprisal model underlies the N400 (20%). We next asked which of the three models is most frequent in the population. This can be assessed via the protected exceedance probability, which is protected against the possibility that differences in model probabilities between participants occur by chance (Piray et al., 2018). Figure 4 (right panel) shows that for the pure surprisal model, the probability that it is the most common model in the population is very low (4%). By contrast, for the combined model, the probability of being the most common model in the population is largest (56%). The pure semantic update model has a slightly lower protected exceedance probability of 40% and thus there is a somewhat smaller chance for this model to be the most common model in the population.

**Figure 4.**
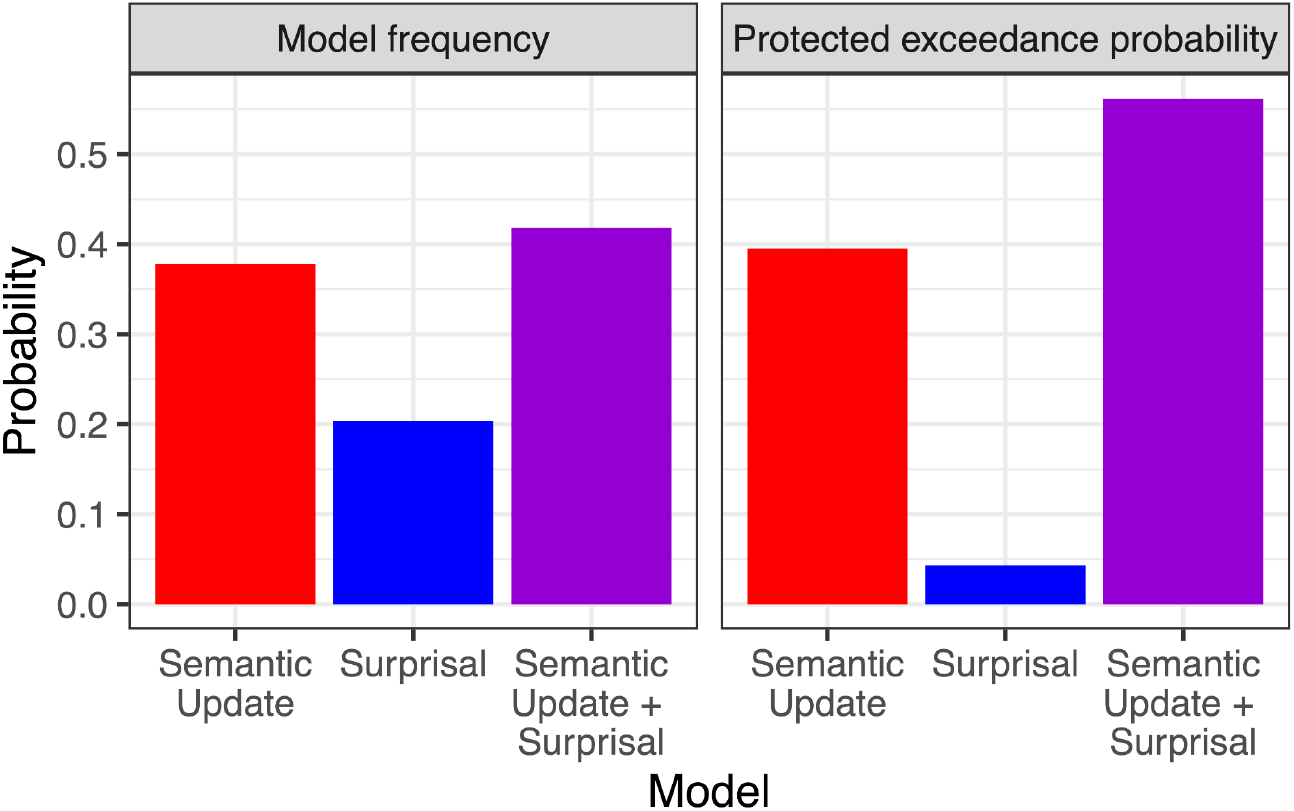
Results from hierarchical Bayesian random-effects model comparison (Piray et al., 2018). Model frequencies (left panel) indicate hierarchical variational Bayesian estimates of how often each model occurs in the population. The combined model (semantic update + surprisal) occurs most often, nearly matched by the pure semantic update model, whereas pure surprisal seems to occur less often. Protected exceedance probability (right panel) indicates the probability that a given model is the most common model in the population. It is protected against the possibility that differences in model probabilities between subjects occur by chance. The results show that a pure surprisal model is not the most common model in the population. Instead, either the combined model or possibly the semantic update model is the most common model.

The hierarchical Bayesian random-effects model also estimates responsibilities, i.e., the probabilities by which each participants’ data was generated by a given model. Comparison of responsibilities (Figure 2b) with parameter estimates from the linear mixed-effects model (Figure 2a) suggest that the eight participants without an individual surprisal effect are classified as purely driven by semantic update (Figure 2b, left panel), that four participants with no clear semantic update effect are purely driven by surprisal (middle panel), and that the remaining ten participants are driven by a combination of both measures (right panel). Interestingly, the six participants with the strongest semantic update effect (the upper six participants in Figure 2) are classified into the pure semantic update model (rather than a combination of both models), which further corroborates the trade-off between computational strategies that was also evident in the negative random-effects correlation of the LMM.

## Discussion

In the current study, we investigated interindividual differences in people’s tendencies to predict words versus sentence meaning, as reflected in N400 amplitudes. This involves investigating random effects correlations in a linear mixed-effects model (LMM) as well as using a random effects model comparison approach suggested by Piray et al. (2018) to test whether in most participants N400 amplitudes are better explained by prediction error at the level of sentence meaning, at the level of the next presented word, or both. Our results from the LMM show that a large semantic update (corresponding to an implicit prediction error at the level of sentence meaning) enhances N400 amplitudes in all participants, whereas large surprisal (corresponding to prediction error at the level of the next presented word) seems to enhance N400 amplitudes in the majority of participants, but that in a subgroup of participants, this effect seems to be reversed. Interestingly, the two effects seem to be negatively correlated, such that participants with a weak or reversed surprisal effect show the strongest influence of semantic update. Moreover, random-effects model selection shows that for large percentages of participants, N400 amplitudes are best explained by (i) pure semantic update or (ii) a combination of semantic update and surprisal, whereas only for few participants N400 amplitudes are best explained by a pure surprisal model.

These results corroborate the notion that N400 amplitudes reflect prediction error both at the word and sentence level, and that surprisal and semantic update effects cannot be reduced to a common factor (Lopopolo & Rabovsky, 2024). Most important for current purpose, our findings suggest that people may differ in their implicit prediction strategies during language comprehension with some participants more focused on predicting sentence (or message) meaning, some participants focused on predicting both sentence meaning and the next upcoming word, and only few participants solely focused on predicting the next upcoming word

The current approach of combining large-scale models implementing different theoretical accounts with advanced model comparison techniques considering differences in computational processes between individuals provides a conceptually novel approach in the language sciences. This approach allows the identification of differences and trade-offs between computational strategies among individuals, as well as the recognition of the most common computational strategies within the population. As a result, it enables more fine-grained investigations of the processes underlying N400 amplitudes, shedding light on the mechanisms involved in language comprehension and prediction in the brain. A limitation of our results is that analysis of protected exceedance probabilities did not allow to unequivocally determine which model is most common in the population, since two models (pure semantic update versus semantic update and surprisal combined) had protected exceedance probabilities considerably larger than zero. On the other hand, we can clearly conclude that surprisal alone is not the most common model in the population.

Intriguing questions for future research will be to see whether the interindividual differences observed here reflect stable differences between people or whether the level of prediction may vary based on situational factors such as for instance attention, cognitive control, or reading strategy. In any case, the observed interindividual differences show a remarkable flexibility of the language system in the brain in terms of the representational level of implicit predictions.

## Competing interests

The authors have no competing interests to declare.

## Acknowledgments

This study was funded via an Emmy Noether grant from the German Research Foundation to Milena Rabovsky (RA 2715/2-1) and via Collaborative Research Center (SFB) 1294 (project B09, grant number 318763901) from the German Research Foundation. The funding agency had no role in the conducted research.

## Notes

### Competing Interest Statement

The authors have declared no competing interest.

## References

Bates, D., Maechler, M., Bolker, B., & Walker, S. (2015). Fitting linear mixed-effects models using lme4. Journal of Statistical Software, 67(1), 1–48. doi: 10.18637/jss.v067.i01

Fitz, H. & Chang, F. (2019). Language ERPs reflect learning through prediction error propagation. Cognitive Psychology, 111, 15–52.

Frank, S. L., Monsalve, I., Thompson, R., & Vigliocco, G. (2013). Reading time data for evaluating broad-coverage models of English sentence processing. Behavior research methods, 45, 1182— 1190.

Frank, S. L., Otten, L. J., Galli, G., & Vigliocco, G. (2015). The ERP response to the amount of information conveyed by words in sentences. Brain and Language, 140, 1–11.

Heilbron, M., Armeni, K., Schoffelen, J.M., Hagoort, P., & de Lange, F. (2022). A hierarchy of linguistic predictions during natural language comprehension. Proceedings of the National Academy of Sciences, 119, 32.

Kuperberg, G.R. & Jaeger, T.F. (2016). What do we mean by prediction in language comprehension? Language, Cognition and Neuroscience, 31(1), 32–59.

Kutas, M. & Hillyard, S.A. (1980). Reading senseless sentences: Brain potentials reflect semantic incongruity. Science, 207(4427), 203–205.

Kutas, M. & Federmeier, K.D. (2011). Thirty years and counting: Finding meaning in the N400 component of the event related brain potential (ERP). Annual Review of Psychology, 62, 621–647.

Kuznetsova, A., Brockhoff, P. B., & Christensen, R. H. B. (2017). lmerTest package: Tests in linear mixed effects models. Journal of Statistical Software, 82(13), 1-26. doi:10.18637/jss.v082.i13

Lopopolo, A. & Rabovsky, M. (2024). Tracking lexical and semantic prediction error underlying the N400 using artificial neural network models of sentence processing. Neurobiology of Language, 5(1), 136–166.

McClelland, J. L., St. John, M. F., & Taraban, R. (1989). Sentence comprehension: A parallel distributed processing approach. Language and Cognitive Processes, 4, 287–335.

Michaelov, J.A., Bardolph, M.D., Van Petten, C.K., Bergen, B.K., Coulson, S. (2023). Strong prediction: Language model surprisal explains multiple N400 effects. Neurobiology of Language.

Palmer, M., Gildea, D., & Kingsbury, P. (2005). The Proposition Bank: An annotated corpus of semantic roles. Computational Linguistics, 31(1), 71–106.

Piray, P., Dezfouli, A., Heskes, T., Frank, M. J., & Daw, N. D. (2019). Hierarchical Bayesian inference for concurrent model fitting and comparison for group studies. PLoS Computational Biology, 15(6), e1007043.

Rabovsky, M., Hansen, S.S., & McClelland, J.L. (2018). Modelling the N400 brain potential as change in a probabilistic representation of meaning. Nature Human Behaviour, 2(9), 693–705.

Rabovsky, M. & McRae, K. (2014). Simulating the N400 ERP component as semantic network error: Insights from a feature based connectionist attractor model of word meaning. Cognition, 132(1), 68–89.

Sayeed, A., Shkadzko, P., & Demberg, V. (2018). Rollenwechsel-english: a large-scale semantic role corpus. European Language Resources Association.

